# Predicting hybrid fitness: the effects of ploidy and complex ancestry

**DOI:** 10.1101/2025.02.13.638087

**Authors:** Hilde Schneemann, John J. Welch

## Abstract

Hybridization between divergent populations places alleles in novel genomic contexts. This can inject adaptive variation – which is useful for breeders and conservationists – or reduce fitness, leading to reproductive isolation. Most theoretical work on hybrids involves haploid or diploid hybrids between two parental lineages, but real-world hybridization is often more complex. We introduce a simple fitness landscape model to predict hybrid fitness with arbitrary ploidy and an arbitrary number of hybridizing lineages. We test our model on published data from maize (*Zea mays*) and rye (*Secale cereale*), including hybrids between multiple inbred lines, both as diploids and synthetic tetraploids. Quantitative predictions for the effects of inbreeding, and the strength of progressive heterosis, are well supported. This suggests that the model captures the important properties of dosage and genetic interactions, and may help to unify theories of heterosis and reproductive isolation.

## 1 Introduction

Hybridization occurs when individuals from genetically differentiated lineages mate and produce offspring. These hybrids carry novel combinations of alleles, exposing allelic effects in different genomic backgrounds (Burch et al., 2024; Peñalba et al., 2024). When alleles function well in their new background, hybridization can facilitate adaptation (e.g. Song et al., 2011; Pardo-Diaz et al., 2012; Abbott et al., 2013; Hedrick, 2013; Kulmuni et al., 2023) – a fact exploited by breeders (East, 1909; Shull, 1909; Gowen, 1952; Suneson, 1956; Gerdes et al., 1999; Gur and Zamir, 2004; Mackay et al., 2020; ter Steeg et al., 2022), and by conservationists (Genovart, 2008; Chan et al., 2019). Conversely, when alleles function poorly, low fitness hybrids may form a barrier to genetic exchange, contributing to reproductive isolation and speciation (Dobzhansky, 1937; Butlin, 1987; Levin, 1985; Hoskin et al., 2005). Hence, studying hybrids and predicting their fitness is important in several areas of biology.

Decades of empirical work has revealed recurrent patterns in hybrid fitness data, suggestive of common and general features of genetic interactions (Kölreuters, 1766; Darwin, 1859; Haldane, 1922; Bateson, 1978; Butlin, 1987; Waser, 1993; Trouve et al., 1998; Turelli and Moyle, 2007; Schilthuizen et al., 2011; Wei and Zhang, 2018; Dagilis et al., 2019; Coyne and Orr, 2004, Ch. 8). Inspired by these data, theoretical work has investigated simple fitness landscape models, to see whether they can generate the patterns observed (Orr, 1995; Barton, 2001; Gavrilets, 2004; Fraïsse et al., 2016; Simon et al., 2018; Satokangas et al., 2020; Schneemann et al., 2024). However, most theory has two major limitations. First, predictions apply only to haploid or diploid hybrids, while empiricists often study higher ploidies (Otto and Whitton, 2000; Birchler, 2013; Washburn and Birchler, 2014; Clo and Kolář, 2022). Second, most predictions apply to hybrids between just two parental lineages, while hybrids can have more complex ancestry – whether in nature (Harvey et al., 2019; Natola et al., 2022; Dean et al., 2024), or in agriculture (Busbice and Wilsie, 1966; Groose et al., 1989; Bingham, 1980; Riddle and Birchler, 2008; Washburn et al., 2019). For example, in the Kerguelen islands, hybrid mussels (*Mytilus edulis* complex) contain ancestry from multiple populations of at least three species (Fraïsse et al., 2021); while on British organic farms, hexaploid winter wheat (*Triticum aestivum*), is grown as Composite Cross Populations, combining 20 inbred lines (Suneson, 1956; Knapp et al., 2020).

Here, we consider a class of fitness landscapes used to study two-parent haploid and diploid hybrids by Barton (2001), Chevin et al. (2014), and De Sanctis et al. (2023). We generalize this model to hybrids of arbitrary ploidy, and with ancestry from an arbitrary number of parental lines. We then compare predictions to two extraordinary datasets, from maize (*Zea mays*; Yao et al., 2020), and rye (*Secale cereale*; Lundqvist, 1966). Both data sets compare diploids and synthetic tetraploids of the same genotypes, and both contain hybrids between multiple inbred lines.

## 2 Model

The aim of this work is to predict the fitness of a hybrid from its genomic composition (Hill, 1982). For diploid hybrids between two parental lineages, A and B, we define the hybrid indices *h*_A_ and *h*_B_ = 1 − *h*_A_, as the proportion of alleles in the hybrid that derive from each parental lineage; and the heterozygosity in ancestry *p*_AB_, as the proportion of loci with one allele from each lineage. Throughout, loci are assumed to segregate independently. Previous work has derived the following prediction.

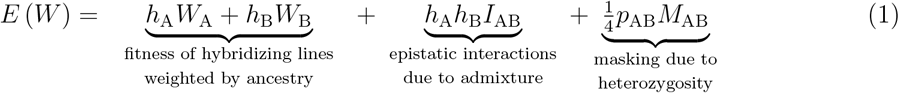

(Chevin et al., 2014; De Sanctis et al., 2023; Schneemann et al., 2024). In this expression, *W* is the fitness of the hybrid, which might have been suitably transformed to give better fit to the model (Lynch and Walsh, 1998, Ch. 10; Fraïsse et al., 2016; Schneemann et al., 2024); *W*_A_ and *W*_B_ are the similarly transformed fitness of the original parental lineages; *I*_AB_ is a measure of pairwise epistatic interactions between the A and B alleles; and *M*_AB_ *>* 0 is a measure of the total amount of fitness evolution that separates them (Chevin et al., 2014; De Sanctis et al., 2023). With this expression, the effects of hybridization partition naturally into (i) the average fitness of the parents, weighted by their contributions to the hybrid; (ii) the effects of admixture, which can vary with the sign of *I*_AB_, and which are maximized for balanced hybrids with *h*_A_ = *h*_B_ = 1*/*2; and (iii) heterotic increases in fitness due to the masking effects of heterozygosity.

Equation 1 was derived from an explicit phenotypic model, of selection on multiple quantitative traits with additive genetics, and fitness declining with the squared Euclidean distance to a phenotypic optimum, such that *W* = −||**z** − **o**||^2^ (Lande, 1976; Turelli, 1985; Orr, 1998; Barton, 2001). This phenotypic model need not hold literally for the fitness landscape to be useful (Martin, 2014; Schneemann et al., 2024), but it does imply strong, testable assumptions about fitness interactions. For example, eq. 1 implies that higher-order interactions are negligible when fitness is suitably transformed (Hill, 1982; Fraïsse et al., 2016), and that deleterious alleles tend to be partially recessive (Manna et al., 2011; Billiard et al., 2021; Schneemann et al., 2022).

Equation 1 applies only to hybrids between a single pair of parental lineages with diploid segregation (see Schneemann et al., 2024 for an application to allotetraploid data). However, it can be easily generalized with a change in notation. To see this, let us redefine *h*_A_ as the proportion of alleles that descend from lineage A at a single locus; so that the hybrid index is its average across loci, now denoted ⟨*h*_A_⟩. It follows immediately that the heterozygosity is *p*_AB_ = 4⟨*h*_A_*h*_B_⟩. Finally, if we number the parental lineages, denoting *P* = 2, then eq. 1 can be written as

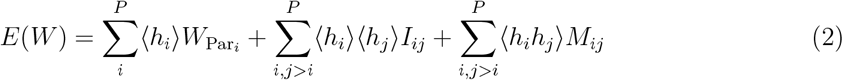

In Appendix S1, we show that eq. 2 applies equally to hybrids with biallelic loci, but containing ancestry from an arbitrary number of parental lineages, *P*, and with arbitrary ploidy, *K*.

### 2.1 The heterozygosity term

Although the three terms in eq. 2 retain their biological meaning, 4⟨*h*_*i*_*h*_*j*_⟩ is not simply heterozygosity in the general case. This is because with high ploidy and complex ancestry, there can be many types of heterozygote, which may contribute differently to masking (Lundqvist, 1966; Ronfort, 1999; Birchler and Veitia, 2012). Accordingly, ⟨*h*_*i*_*h*_*j*_⟩ is a weighted sum of the proportions of the different types of ancestry heterozygote. For example, with a two-parent tetraploid hybrid (*P* = 2, *K* = 4), two types of heterozygote are possible: balanced (AABB) and unbalanced (AAAB or ABBB), for which 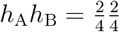 and 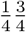, respectively. It follows that

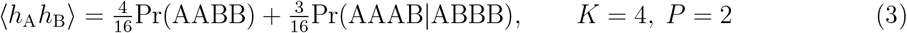

Adding a third parental lineage, C, adds two new types of heterozygote with A and B ancestry, and so

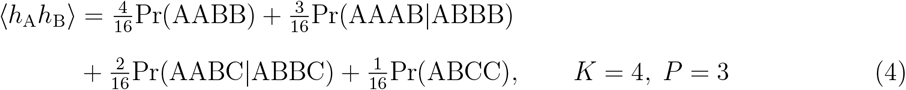

and in general:

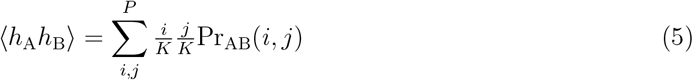

where Pr_AB_(*i, j*) is the proportion of loci with *i* alleles of A ancestry and *j* alleles of B ancestry. Let use also note here that the sum 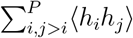 is closely related to Wright’s inbreeding coefficient, *F* (Wright, 1922): the probability that two alleles chosen without replacement from the same locus, are identical by descent (i.e. descend from the same parental lineage). The two quantities are related via

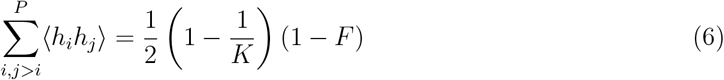

## 3 Inbreeding depression with selfing

One of the major differences between ploidies is the rate at which they lose heterozygosity under inbreeding (Muller, 1914; Haldane, 1930). Given the relationship between our measure of heterozygosity and Wright’s inbreeding coefficient *F*, eq. 2 can be considered as a multilocus generalization of single-locus inbreeding theory, including epistatic interactions. We can also use existing results for changes in *F*. For example, with selfing we have

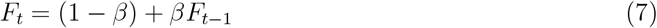

(Wright, 1969; Crow and Kimura, 1970) where the rate of change with generation *t* depends on the ploidy, via

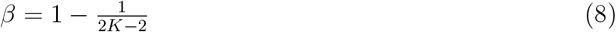

(Hardy, 2015), assuming random chromosome segregation during meiosis without double reduction (Gallais, 2003). Combining these results with eq. 6 we have

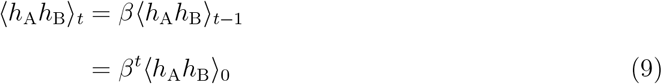

We can now consider a first-generation hybrid between lines A and B, subject to selfing. In this case, any loss of fitness must be due to loss of heterozygosity (the third term of eq. 2), and so we have

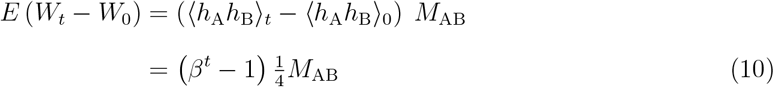

where *β* given by eq. 8, and we have used the fact that ⟨*h*_A_*h*_B_⟩_0_ = 1*/*4 in the initial F1.

### 3.1 Testing the predictions in maize

To test the prediction of eq. 10 we used data from Yao et al. (2020). These data comprise hybrids of four inbred lines of maize (*Zea mays* lines A188, Oh43, W22 and B73), here labelled A, B, C and D. In the main text, we consider only the A×B and C×D crosses, which were generated in both diploid and tetraploid form, via nitrous oxide gas treatment of the diploid zygotes (Kato and Birchler, 2006; Yao et al., 2020). Appendix S2 discusses attempts to interpret the AB×CD cross and crosses only available as diploids.

Yao et al. selfed these initial F1 for seven generations (with measurements at generations 0, 1, 3, 5, and 7 in five replicate experiments). We chose ear length as the available trait closest to overall plant fitness or productivity. We then Box-Cox transformed the measurements to minimize the skewness of the data around their means for each generation (see Appendix S2 for full details), and fit eq. 10 using standard least-squares non-linear regression (Baty et al., 2015). The model includes two parameters (*β* and *M*_*ij*_), but the cross- and ploidy-specific normalization of the data by Yao et al., 2020 means that only the *β* parameter is readily interpretable.

Figure 1A shows the results of the fitting. As expected, the tetraploids lose fitness at a slower rate, matching the predictions of eq. 8 that *β* = 1*/*2 for diploids and *β* = 5*/*6 for tetraploids (Fig. 1B). With the tetraploid data, we can go further, and estimate the weights in eq. 3, which determine the effects of dosage, via the relative contributions of balanced and unbalanced heterozygotes. In this case, we fit the model

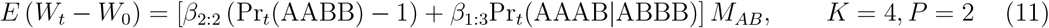

**Figure 1:**
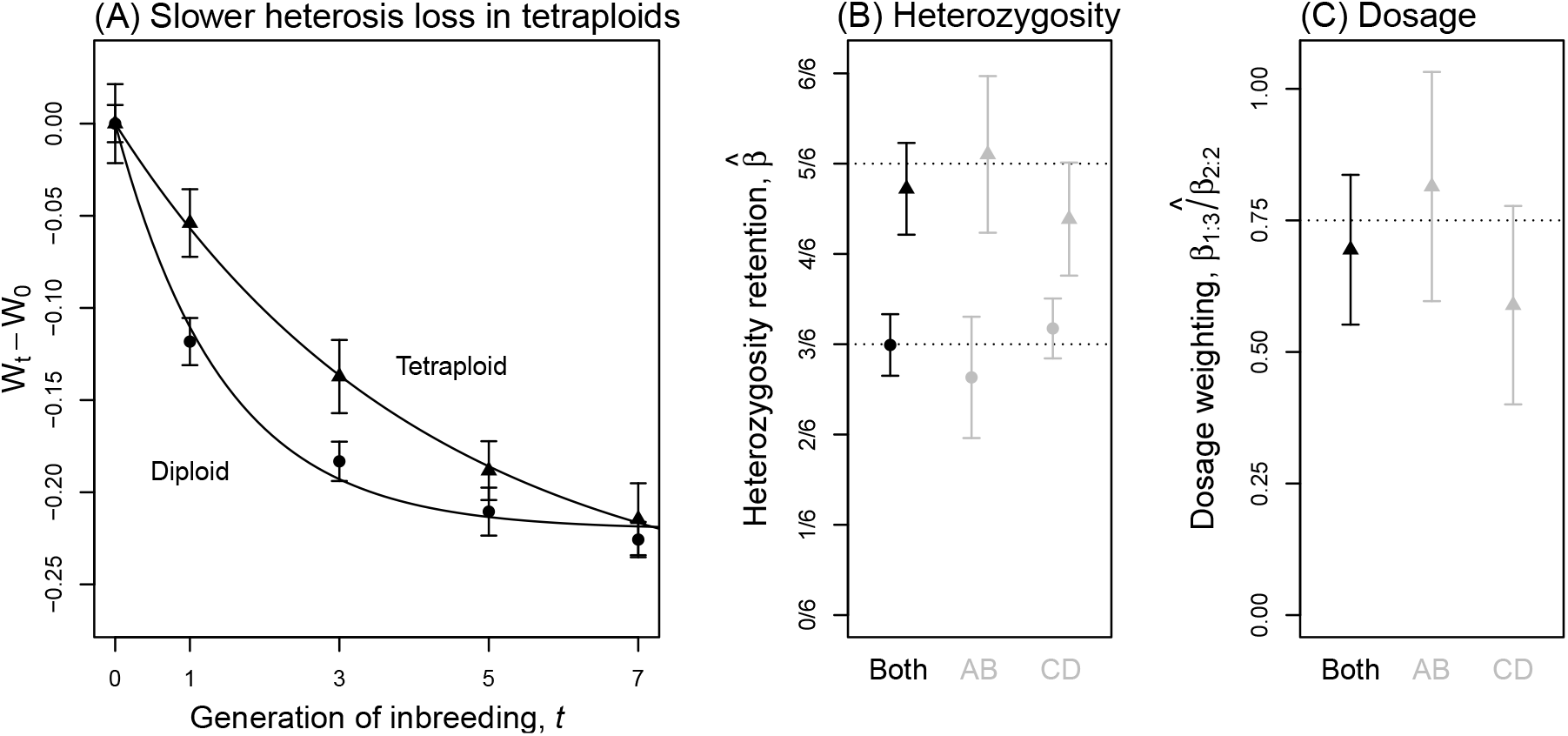
Reanalysis of hybrid maize data from Yao et al. (2020). **(A)** The decline in fitness over 7 generations of inbreeding is slower in tetraploid (▴) than in matched diploid (•) maize crosses. Points and bars show means and standard errors for ear length, transformed to minimize the skew around the within-generation means (see Appendix S2). Lines show the least-squares fit of the non-linear model of eq. 33 (Baty et al., 2015) with *β* fixed at its expected value. **(B)** Estimates of the parameter *β*, which captures the retention of heterozygosity, with 95% confidence intervals (Baty et al., 2015). Estimates of *β* match predictions of *β* = 1*/*2 for diploids, and *β* = 5*/*6 for tetraploids (eq. 8). **(C)** Fitting eq. 11, estimates the different effects of balanced and unbalanced heterozygotes in tetraploids, and agrees with predictions of *β*_13_*/β*_22_ = 3*/*4 (confidence intervals on the ratio used the “delta method”; Fox et al., 2001; Fox and Weisberg, 2019).

With

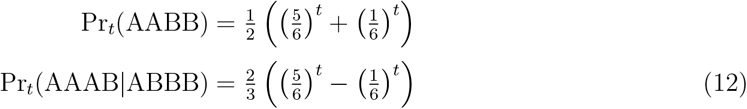

(which follow from recursions in Haldane, 1930). Then, eq. 3 implies the prediction 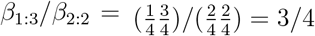. Results, shown in Figure 1C, suggest good agreement with this prediction.

## 4 Progressive heterosis with multiple parents

We can further test the predictions of eq. 2 by investigating the fitness of hybrids with ancestry from more than two parental lineages. Here we use a classic dataset from Lundqvist (1966), involving six inbred parental lines of diploid rye (*Secale cereale*; steel variety, lines 19-23 and 25), here labelled A-F. Unlike the modern data of Yao et al. (2020), Lundqvist (1966) studied only two generations of hybridization, and reported only means for each cross; but the dataset includes not only the 6 parental lines and the 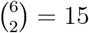 possible F1, but also the 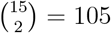 possible F2. These data allow us to test the effects of complex ancestry, because the F2 had ancestry from either *P* = 2, 3 or 4 parents, depending on the number of parents shared by the crossed F1. Moreover, we can also compare results for diploids and tetraploids, since colchicine treatment was used to generate synthetic tetraploids of the six inbred lines; and of the 126 crosses, only a single F1 was absent in tetraploid form.

Figure 2A shows a cartoon of all of the cross types present in the Lundqvist (1966) data, along with predictions from eq. 2, assuming identical fitness for the diploid and tetraploid F1. Three main patterns are evident in Figure 2A. First, there is a large increase in fitness between the parents and F1. As shown by Hill (1982) this F1 heterosis can involve both dominance and epistasis, and if we average over all F1, we have

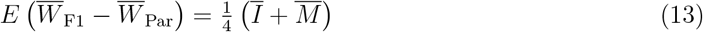

where Ī and 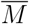 are averages over all pairs of parental lineages. It follows that heterosis will be positive unless the epistasis, *Ī*, is strongly negative; and heterosis will differ among ploidies only if *Ī* and 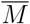 also differ.

**Figure 2:**
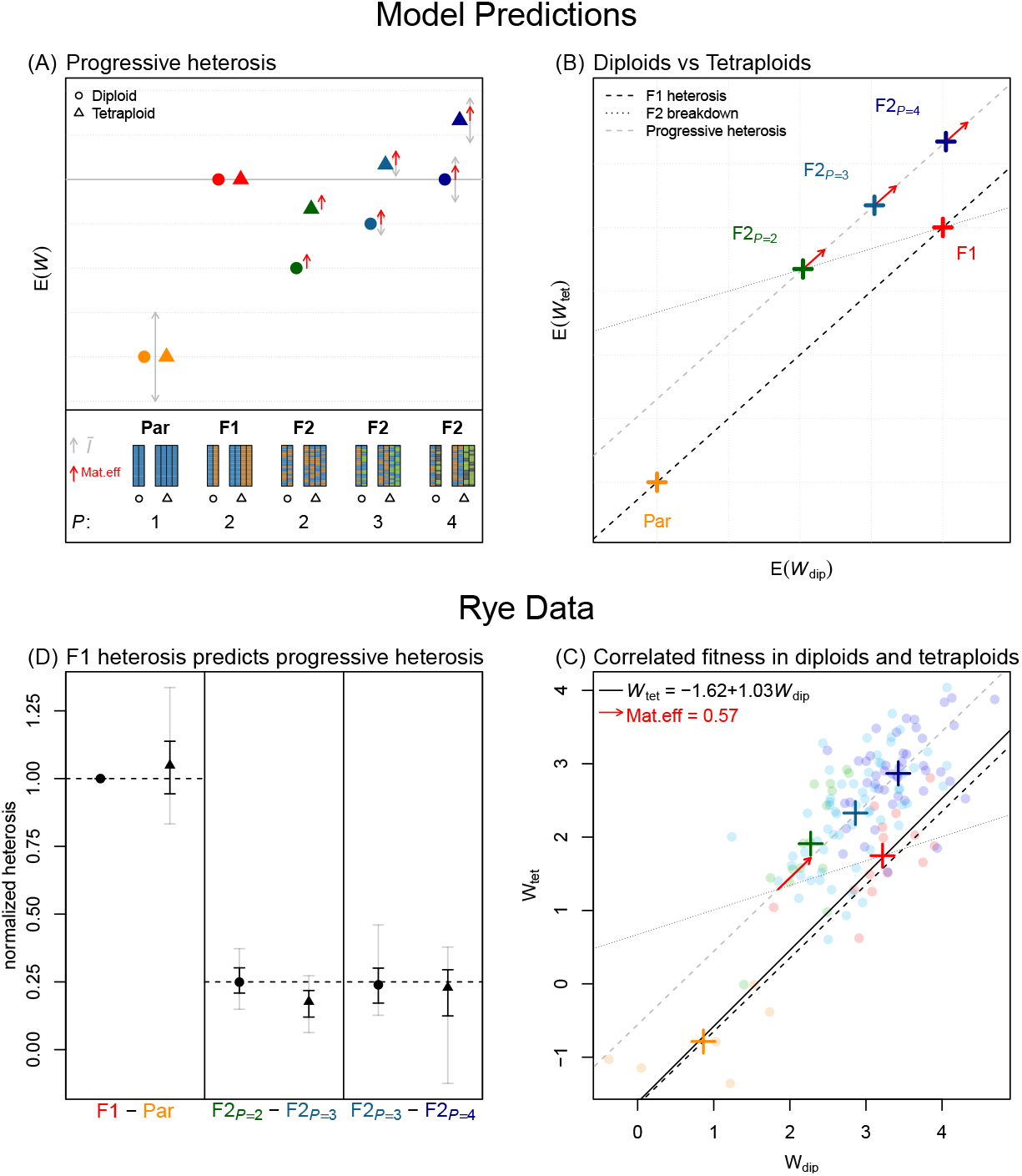
Reanalysis of hybrid rye data from Lundqvist (1966). **(A)** Predicted fitness for different crosses in diploids (•) and tetraploids (▴), with ancestry from *P* = 1, 2, 3, or 4 parental lines (eq. 2). Ploidies differ only in their F2 breakdown (fitness reduction between F1 and F2; eq. 14). Fitness epistasis, *I* (gray arrows), can vary in sign, and affect both F1 heterosis (improvement between Parents and F1; eq. 13) and progressive heterosis (improvement from adding parents to the F2; eq. 18). A maternal affect (red arrows) may increase the fitness of all F2, because they were grown from fitter F1 mothers. **(B)** Slopes that indicate F1 heterosis and progressive heterosis should match (eq. 18); while the F2 breakdown slope should be 1/3 smaller (eq. 14), but maternal effects make the latter difficult to test. **(C)** Rye data, with total kernel yield transformed to maximize the correlation between ploidies for the Parent and F1 measurements (see Appendix S3), and individual crosses (•) compared to means of cross types (+). The best-fit SMA regression (Warton et al., 2012) has a slope close to 1 (solid line), implying that polyploidization little changed the *M*_*ij*_ and *I*_*ij*_ parameters. **(D)** F1 heterosis in diploids accurately predicts the progressive heterosis, in both diploids and tetraploids (eq. 18). Results show the difference in *W* for the crosses indicated, normalized by the difference between diploid Parents and F1. Error bars are jackknife resamples, removing any cross with ancestry from each of 6 inbred lines (black bars), or each of 15 pairs of lines (gray bars).

The second pattern in Figure 2A is the loss of fitness between the F1 and F2. The strength of F2 breakdown does differ between ploidies, because – as discussed above – they lose heterozygosity at different rates. Using eqs. 8 and 10 we have

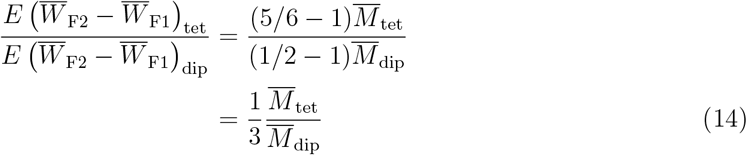

The final pattern in Figure 2A is the steady increase in F2 fitness with the number of parental lineages, *P*. Sometimes, F2 fitness exceeds the F1, a phenomenon known as progressive heterosis (Washburn and Birchler, 2014). To understand this, let us first consider the average fitness of all F2 derived from *P* parental lineages:

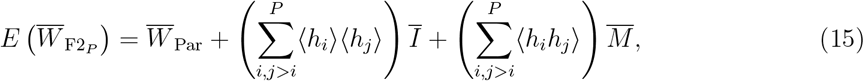

where the sums in brackets would be identical for any *P*-parent F2. It is quick to calculate the first sum as 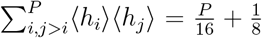, while the second sum follows from the F2 inbreeding coefficient, which is

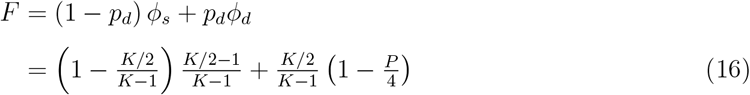

where *p*_*d*_ is the probability that two alleles, drawn at random from an F2 locus, derive from different F1 gametes; and *ϕ*_*s*_ (*ϕ*_*d*_) is the probability that alleles drawn from the same gamete (different gametes) are identical by descent. Substituting this result into eq. 6 gives us 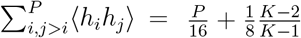. We can now derive the conditions for progressive heterosis, by comparing the fitness of the F1 and the four-parent F2.

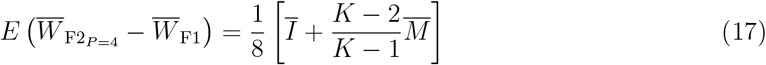

So in diploids (*K* = 2) progressive heterosis will appear only when epistasis is positive (*Ī >* 0); while in tetraploids (*K* = 4) progressive heterosis will always appear, unless epistasis is very strong and negative, such that 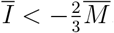. Finally, we can relate the strength of the progressive heterosis to the F1 heterosis, by noting that

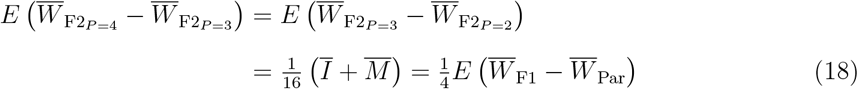

So for any ploidy, the fitness gain from adding an additional parent to the F2, is a quarter of the original F1 heterosis. Note that the first line of eq. 18 – the linear change with *P* – holds quite generally, because the total number of heterozygous loci changes linearly for both ploidies (Lundqvist, 1966). However, the second line – which relates F1 heterosis and progressive heterosis – only holds in polyploids with our particular weighting of the different types of heterozygote (eq. 5). All three predictions (eqs. 13, 14 and 18) are summarized in Figure 2B.

### 4.1 Testing the predictions in rye

To test these predictions in the rye data, we used total kernel yield as the best-available proxy for plant fitness. We then Box-Cox transformed the data to maximize the fit (*r*^2^) of a Standardized Major Axis regression between the diploid and tetraploid measurements (Warton et al., 2012), but using only the fixed genotypes – i.e. the 6 parents and 14 available F1. We then used this best-fit regression to impute a value for the missing tetraploid F1 (see Appendix S3 for full details). The resulting data are plotted in Figure 2C.

The first notable result in Figure 2C is that the best-fit regression line between the diploid and tetraploid fixed genotypes (solid line), is very close to a 1:1 line, but with a non-zero intercept (dashed line). This is consistent with polyploidization inducing a constant fitness cost for all genotypes, but with the *M*_*ij*_ and *I*_*ij*_ parameters remaining constant. This is supported by model selection using the complete data set of 252 crosses (see Appendix S3 for full details).

The second observation from Figure 2C is the failure of the prediction of hybrid breakdown (eq. 14). In fact, the two-parent tetraploid F2 are generally fitter than the equivalent F1 (see also Figure S4). Lundqvist (1966) also remarked on this surprising aspect of his data, and speculated that a strong maternal effect might have provided a fitness boost to all F2 plants (who came from high-fitness F1 mothers), relative to the F1 and parental plants (which came from low-fitness inbred mothers). Red arrows in Figure 2A-B show how a maternal effect would affect our predictions. Note that an effect might apply equally across ploidies, and yet remove hybrid breakdown completely only in the tetraploids (as observed). We could not test directly for a maternal effect, since crosses were not generally made in both directions (Lundqvist, 1966). However, given our other assumptions, models with a maternal effect were preferred (see Appendix S3).

Finally, in Figure 2D, we test the prediction of eq. 18. Given the apparent constancy of the Ī and 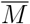 parameters, we used the observed heterosis in diploid F1, to predict the effects of adding parents to F2 of both ploidies. While tetraploid two-parent F2 were slightly fitter than predicted, the quantitative agreement was otherwise good. For example, the strength of F1 heterosis in diploids well predicted the benefit of adding a fourth parent to tetraploids.

## 5 Discussion

We have extended existing models for predicting hybrid fitness (Barton, 2001; Chevin et al., 2014; De Sanctis et al., 2023), so that they apply to hybrids of arbitrary ploidy, and with an arbitrary mix of ancestry. While previous diploid predictions can be applied to allopolyploids, where segregation is effectively diploid (see the reanalysis of *Brassica* data from Hauser et al., 1998 by Schneemann et al., 2024), eq. 2 applies to any mode of segregation. This main result is comparable to previous work on the quantitative genetics of autopolyploids (e.g. Fisher et al., 1965; Busbice and Wilsie, 1966; Bennett, 1976; Gallais, 2003), but reduces the number of parameters by assuming an underlying phenotypic model and biallelic loci. This model allows for pairwise epistatic effects (e.g. Gallais, 2003, p.183) – a feature missing from some earlier single-locus analyses (e.g. Wright, 1977, pp. 21-25) – but it does entail strong assumptions about dominance, and especially the masking effects of different types of heterozygote (e.g. AAAB vs. AABB; eq. 5). While a range of different assumptions have been explored in the breeding literature (e.g. Lundqvist, 1966; Ronfort, 1999), our assumptions gave a good fit to the data analyzed here (e.g. Figs. 1B-C and 2D).

### Explaining heterosis

We applied our model to two published data sets (Lundqvist, 1966; Yao et al., 2020), both of which show heterosis, i.e. increased fitness for early generation hybrids (Kölreuters, 1766; Darwin, 1859; Gowen, 1952; Birchler, 2013; Muraro et al., 2022). There are longstanding debates about the causes of heterosis – and especially the adequacy of the simplest theory: that the parental lines fixed recessive deleterious alleles, whose effects are masked in the hybrids (Bruce, 1910; Keeble and Pellew, 1910; Crow, 1948; Lippman and Zamir, 2007; Birchler, 2013; Washburn and Birchler, 2014). The same process of masking – sometime called “the dominance theory” or “the complementation theory” – could, in principle, explain the progressive heterosis observed in polyploids with multiparent ancestry (Figure 2A-B; Jones, 1918; Stringfield et al., 1950; Bingham, 1980; Groose et al., 1989). Our fitness landscape model offers a generalization of the classical masking theory, and may help to resolve some of these debates.

For example, if heterosis is caused by masking of deleterious alleles, then we could expect to select out these alleles, thereby “fixing” the heterosis – but this has not always proven possible (Jones, 1917; Vetukhiv, 1954; Birchler, 2003; Koltunow and Tucker, 2003; Washburn et al., 2019). Here, we have shown that F1 heterosis (eq. 13), and progressive heterosis (eq. 17) can both arise via masking, even when there are epistatic fitness interactions between the parental alleles. Nevertheless, with epistasis, “deleterious” alleles might be deleterious only in some genetic backgrounds (Hwang et al., 2017; Xie et al., 2022), and this could make them difficult to purge. Thus, our model may help to reconcile the observation of heterosis arising from masking, with the difficulty of fixing heterosis through selection on hybrid populations (see Schneemann and Welch, 2025).

More directly, the magnificent data of Yao et al. (2020) were originally reported as evidence against the masking theory, because no evidence was found that diploids and tetraploids lost fitness at different rates. Here, by contrast, we did find a difference in the predicted direction (Figure 1B). One explanation is that the form of the relationship is non-linear (eq. 33), and so unlikely to be detected by fitting a standard linear model (Yao et al., 2020), especially if data are not transformed to meet the assumptions of the model fitting (see Appendix S2).

Non-linearity and scales of measurement are relevant for a second debate about the masking theory. If deleterious mutations are successfully removed, then the theory predicts a reduction in the potential for heterosis, which is not always observed (Duvick et al., 2005; Troyer and Wellin, 2009). However, the quantitative change in the heterosis with selective improvement of the parents will depend on the data transform used. So, for example, yield might show increasing heterosis, while log yield shows decreasing heterosis. As such, observed changes in heterosis (Duvick et al., 2005; Troyer and Wellin, 2009) are difficult to use as evidence either for or against the masking theory (Birchler, 2013; Washburn et al., 2019; see Appendix S4 for more details). Whatever the outcome of these debates, the fitness landscape model used here, incorporating, as it does, different types of gene action (Manna et al., 2011; Sellis et al., 2011) may facilitate quantitative analysis of hybrid fitness data and aid progress towards a unifying theory of heterosis (Birchler, 2013).

### Polyploidization and hybridization

While both reanalyses presented here (Figures 1–2) concern heterosis, we hope the theoretical results will apply more widely. If hybridization can contribute to biodiversity by injecting adaptive variation (Kulmuni et al., 2023; Peñalba et al., 2024), it can also do so by failing – thereby increasing reproductive isolation. Polyploidization may play a special role in both the positive and negative processes. For example, polyploid hybrids could have enhanced adaptive potential – not only by better masking their ancestors’ deleterious mutations (as discussed above), but also by combining their adaptations in a single individual, or by releasing pleiotropic constraints via specialization of the subgenomes. On the other hand, ploidy differences can pose instant and strong barriers to gene exchange (Marks, 1966; Otto and Whitton, 2000; Turelli et al., 2001; Hegarty and Hiscock, 2004; Soltis et al., 2014), and can themselves be caused by hybridization (Masterson, 1994; Lewis, 2012). Indeed, these are all possible explanations for the abundance of polyploids in nature.

In our approach, the positive and negative effects of hybridization are governed by the size and magnitude of the parameters (i.e. the *M*_*ij*_ and *I*_*ij*_), and the underlying phenotypic approach allows us to predict their changes under different modes of evolutionary divergence (Chevin et al., 2014; Simon et al., 2018; Schneemann et al., 2020; De Sanctis et al., 2023; Schneemann et al., 2024). The phenotypic model also makes a naive prediction about how these quantities will change with polyploidization itself. Since *M*_*ij*_ and *I*_*ij*_ are both proportional to *K*^2^ (see Appendix S1 eqs. 30-32), then – all else being equal – tetraploidization of diploids should increase both quantities by a factor 4^2^*/*2^2^ = 4. This naive prediction could not be rigorously tested by either data set analyzed here, although the the rye data gave suggestive and surprising evidence that similar *M*_*ij*_ and *I*_*ij*_ applied for both ploidies. While it seems unlikely that any simple fitness landscape model could capture all of the effects of polypoidization – with its diverse morphological, cytological and genetic consequences, including genome instability (Otto and Whitton, 2000; Riddle et al., 2006; Soltis et al., 2014), we hope that our predictions for the masking and dosage effects in polyploids will help to tease these factors apart.

## Acknowledgments

We are grateful to James Birchler, Sanvesh Srivastava, and Rebecca Doerge for providing data and clarification. We are also grateful to Roger Butlin and Andrea Manica for helpful comments on an early draft, and to Nicolas Bierne for encouragement. HS acknowledges support from the Wellcome Trust program in Mathematical Genomics and Medicine (RG92770), and from the European Union’s Horizon 2020 research and innovation programme under the Marie Sklodowska-

Curie Grant Agreement No.101034413. The authors distribute this work under a CC BY 4.0 copyright license.

## S1 Appendix 1

### Derivation of main result

In this Appendix we derive eq. 2, starting with an explicit model of selection on phenotypes. The approach follows Lande (1981), Chevin et al. (2014) and De Sanctis et al. (2023), but is generalized to arbitrary ploidy, *K*, and hybrids between an arbitrary number, *P*, of parental lineages. The model posits that selection acts on several quantitative traits, such that the transformed fitness of any individual is given by

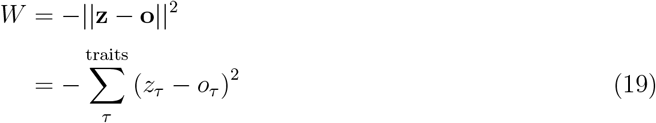

where *z*_*τ*_ is the individual’s value of trait *τ*, and *o*_*τ*_ is this trait’s optimal value. If fitness is log transformed, so that *W* denotes log fitness, then this is a standard model in quantitative genetics (e.g. Lande, 1976). From eq. 19, it follows immediately that the expected transformed fitness for any group of individuals, is given by

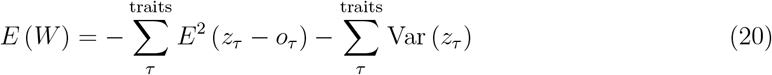

Let us note immediately that eq. 20. involves a simple sum over traits, and so below we calculate quantities for a single trait, and drop the *τ* subscript for brevity. Let us now consider hybrids between *P* parental populations, characterized by genetic variation at several loci. Our major simplifying assumptions are that (1) all variable loci are biallelic; (2) the genetics of the traits are entirely additive; and (3) there are no statistical associations between alleles within the parental populations (an assumption that holds automatically when the parents are inbred lines, with no within-population variation). We now designate one allele at each locus as the “focal allele”, and note that the number of focal alleles carried at each locus will be binomially distributed between 0 and *K*, so that the mean and variance of a trait in parental population *i* = 1, 2, …, *P* are

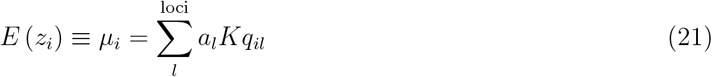

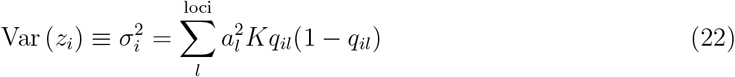

where *q*_*il*_ is the frequency of the focal allele at locus *l* in population *i*, and *a*_*l*_ is the (additive) phenotypic effect of the focal allele on the trait (note that, without loss of generality, we can neglect the trait value of the reference genotype, containing no focal alleles, in the derivation).

Let us now turn to hybrids between these populations. Consider, first, a single locus in the hybrid. The number of focal alleles carried at this locus will again vary between 0 and *K*, and will also depend on the parental allele frequencies. For the hybrid, however, we must also consider the proportions of ancestry from each population at the locus. Following the main text, we denote these proportions as *h*_1_, *h*_2_, …, *h*_*P*_. Then, denoting the number of focal alleles as *X* (and dropping the locus subscript for brevity), we have

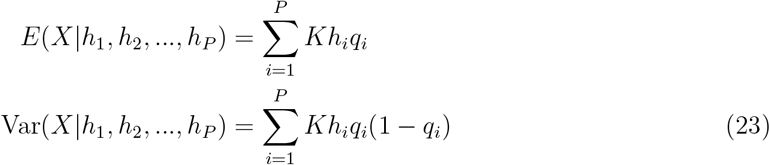

where all subscripts refer to the parental population. Next, we treat the *h*_*i*_ themselves as random variables, which may vary between loci and hybrid individuals. Following the main text, we denote their mean and covariance as ⟨*h*_*i*_⟩ and ⟨*h*_*i*_*h*_*j*_⟩−⟨*h*_*i*_⟩⟨*h*_*j*_⟩, respectively. Our derivation then makes the crucial assumption that the distribution of ancestries is the same at all loci. With this assumption, it follows that the unconditional mean and variance of *X* are

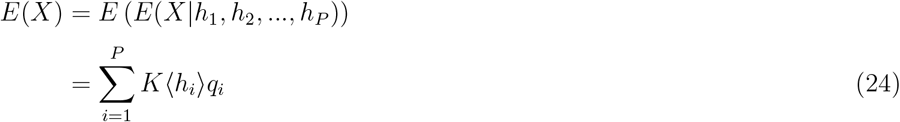

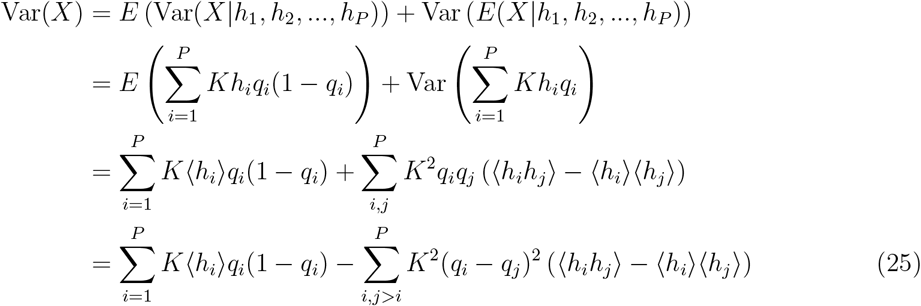

where the last line uses 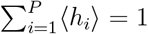, which follows because all ancestry in the hybrid must come from one of the *P* parental lines.

Finally, we can apply these results to calculate the two terms in eq. 20 for a collection of hybrids. To do this, we sum over loci, but still consider only a single trait. For the first term, we note from eq. 21 that the hybrid trait mean is simply

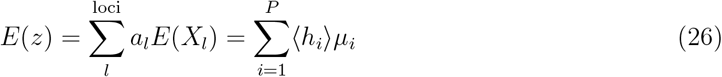

and so

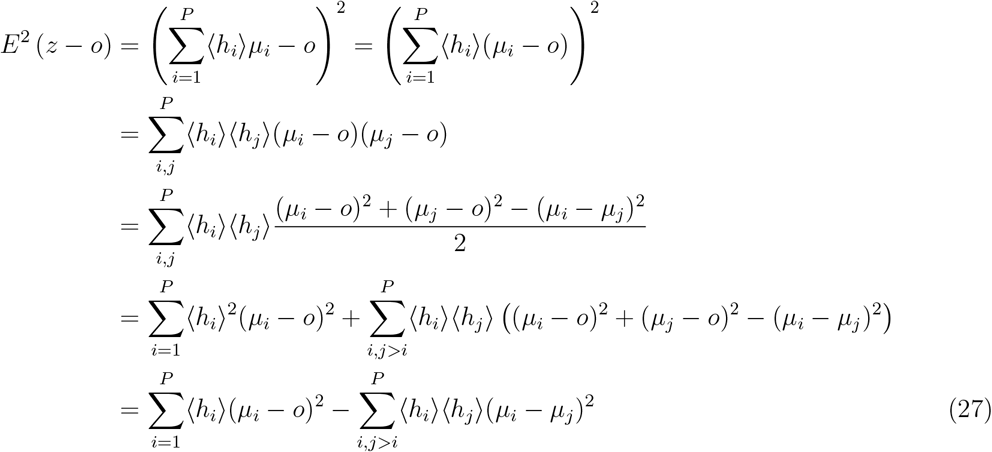

Then, for the variance, using eq. 22 we have

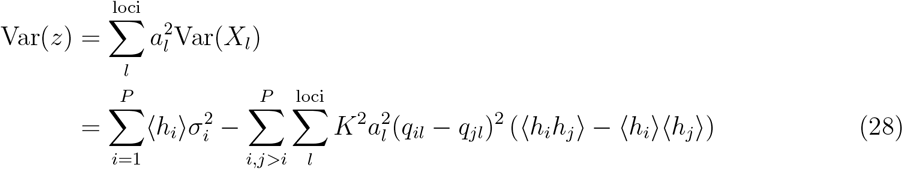

The derivation of eq. 2 is completed by noting that, from eq. 20, the mean transformed fitness of parental line *i* is

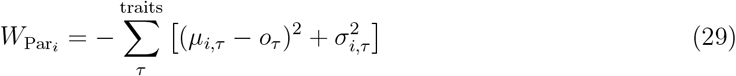

and then defining

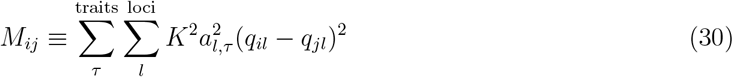

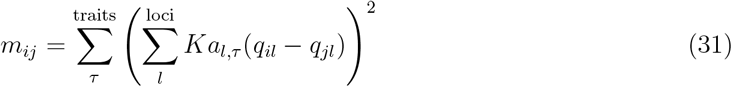

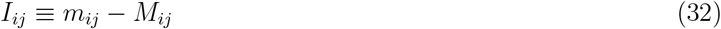

For the interpretation of these quantities, and especially the identification of *I*_*ij*_ with fitness epistasis, see Chevin et al. (2014), Schneemann et al. (2020), De Sanctis et al. (2023) and Schneemann et al. (2024).

## S2 Appendix 2

### Details of reanalysis of Yao et al. (2020) maize data

#### S2.1 Data description

The dataset from Yao et al. (2020) comprises hybrids of four inbred lines of maize (*Zea mays* lines A188, Oh43, W22, and B73), here labelled A, B, C and D. For each cross, seven generations of selfed offspring are available, with three kernels taken from different ears after one round of selfing, such that subsequent generations include three selfing lineages. All generations of plants were grown simultaneously in a randomized complete block design (across two fields in 2008 and three fields in 2009) and yield-related traits were recorded after 1, 3, 5 and 7 generations of selfing. Yao et al. (2020) reported data for 9 distinct yield-related traits (ear length, 4-week, 6-week and adult height; leaf length and width; tassel branch number; and silk and flower times). Before carrying out any analyses, we chose ear length as the trait closest to overall plant productivity or fitness. On average 45 (min. 3, max. 76) individuals were recorded per year, field, ploidy, cross, and generation combination. The data reported are individual measurements normalized by the value of the corresponding F1 (same cross, ploidy and year), and measurements of the inbred lines themselves were not included. For both reasons, we could not use these data to estimate *M*_*ij*_ or investigate the effect of polyploidization *per se*.

#### S2.2 Model fitting

The maize data showed a large degree of skew in the residuals around their means, violating the assumptions of least-squares model fitting. As such, we used a Box-Cox transformation, replacing the raw measurement *x* with (*x*^*λ*^ − 1)*/λ*, and choosing the value of *λ* that minimized the absolute skew (third central moment) averaged over each generation of the data. Figure S1A-B show results for the crosses used in the main text (A×B and C×D); these results show the absolute skew of the transformed data with 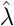 is negligible, but increases rapidly for higher or lower *λ* values. After transformation, we fit our non-linear model to these data (eq. 10), using the least-squares method implemented in the *R* function *nlsLM* (Elzhov et al., 2022; R Core Team, 2021) and calculated confidence intervals with the asymptotic method implemented in the function *confint2* (Fox and Weisberg, 2019). To calculate the dosage weighting ratio *β*_1:3_*/β*_2:2_ in Figure 1E, we used the *R* function *deltaFunction* (Fox and Weisberg, 2019) to obtain the maximum likelihood estimate and confidence intervals on the ratio.

**Figure S1:**
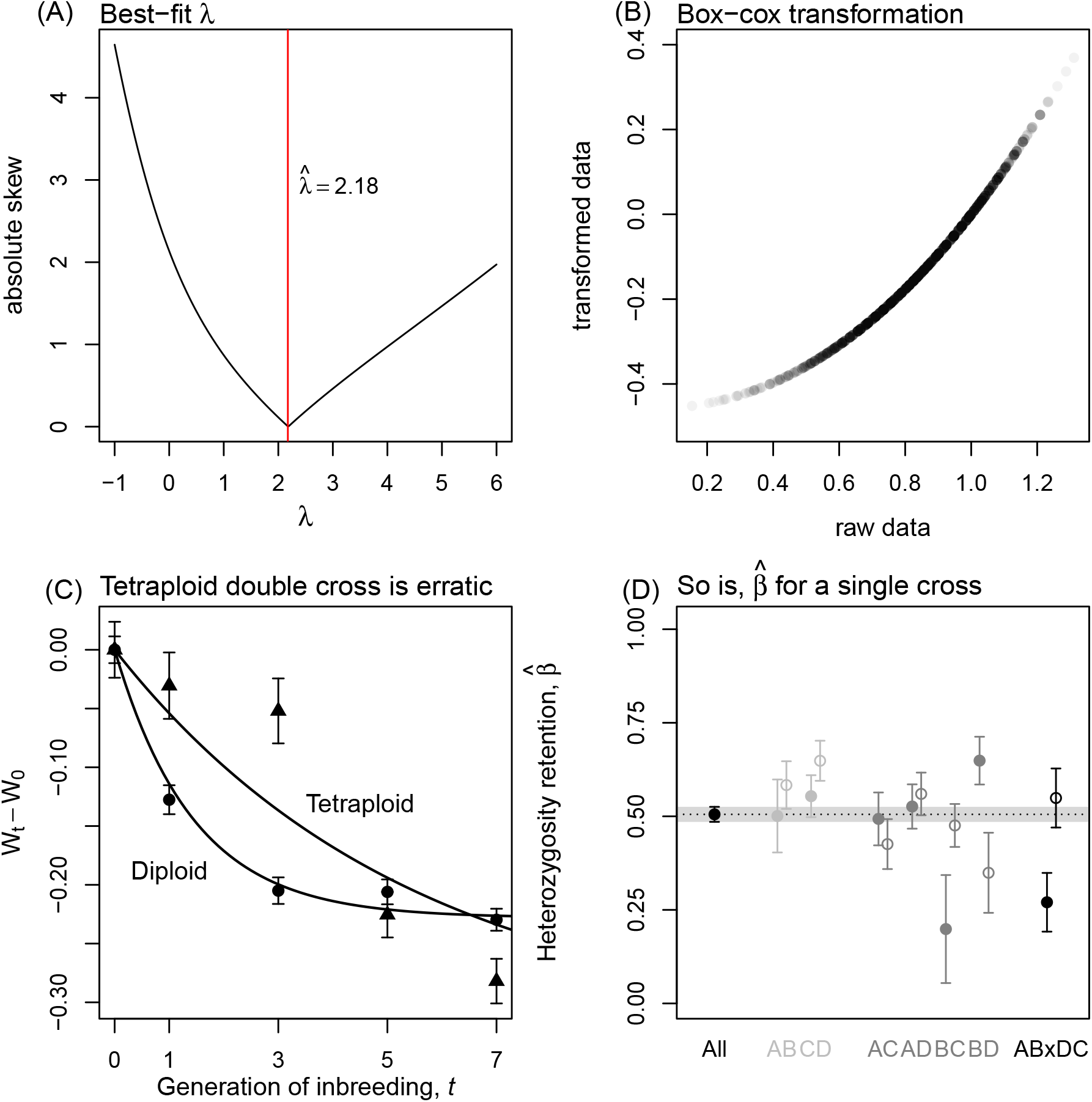
Maize data transformation and cross direction effect. **(A)** The best-fit *λ* used in the Box-Cox transformation of the maize data minimizes mean absolute skew per generation. **(B)** The corresponding Box-Cox transformation. **(C)** The diploid 4-parent cross (•) fits the model predictions, while the tetraploid (▲) fits poorly and shows an erratic loss of heterosis. Plotting details match Fig. 1A. **(D)** Even for diploid data, estimates of *β* are highly heterogeneous between crosses and cross directions, although the estimate from all diploid data combined very closely fits the prediction of *β* = 1*/*2 (dotted line). The filled and open points show estimates for the cross listed on the x-axis and its reciprocal respectively.

#### S2.3 Predictions for four-parent data

Yao et al., 2020 also provided data for other crosses: a four-parent double cross (AB×CD) in both diploid and tetraploid state, and all six of the possible two-parent crosses in diploid state. The diploid crosses were available in both cross directions, while all of the tetraploid crosses (including those in the main text) were present in just one cross direction.

The prediction in eq. 10 generalizes readily to crosses with more than two parents if we weight the *M*_*ij*_ according to their contribution to the F2 (i.e. weighted by ⟨*h*_*i*_*h*_*j*_⟩_0_, see Table S1). For example, in the diploid double cross four different types of heterozygote are expected at equal frequency (AC, AD, BC and BD), while in the tetraploid, all 6 types of heterozygotes are present. As such, we have

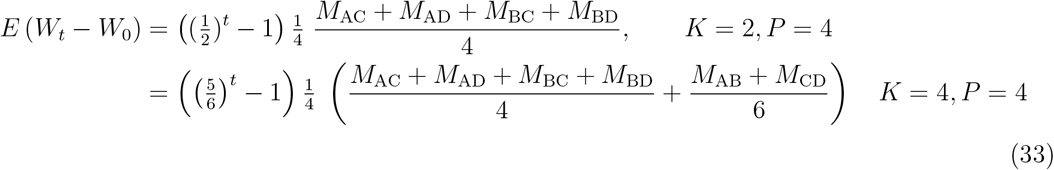

Thus, predictions for both ploidies can be written in the form 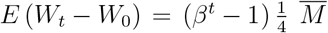 which we can fit to the data.

Figure S1C shows this prediction fit to the four-parent data (diploid: circles; and tetraploid: triangles), Box-Cox transformed with 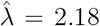 as above. Results show that the diploid points, which are averaged over the two cross directions, AB×DC and DC×AB fit the prediction fairly well. By contrast, the tetraploid 4-parent cross, which was available in just one cross direction (BA× CD) fits very poorly. In fact, this cross shows an erratic patterns of loss of heterosis, with little change between generations 1 and 3, but a steep drop in fitness between 3 and 5.

This result might be a true failure of the model, or simply noise in the data. A general impression from the maize analyses is that, even with this large dataset, patterns in the data are relatively noisy unless we average across crosses and/or cross directions (see Figure 1B-C), which were not available in this case. For comparison, Figure S1D shows *β* estimates for each diploid cross (indicated along the x-axis) and direction (filled vs open points). Only when we average over these crosses do we find that our estimate of *β* matches the prediction of 1*/*2 very closely, while individual estimates can deviate substantially from this value. This prediction of *β* = 1*/*2 stems purely from Mendelian segregation, rather than any assumptions of the fitness landscape, suggesting that the deviation of the tetraploid 4-parent from model predictions might similarly be a result of sampling noise.

## S3 Appendix 3

### Details of reanalysis of Lundqvist (1966) rye data

#### S3.1 Data description

The dataset from Lundqvist (1966) involves six inbred parental lines of diploid rye (*Secale cereale*; steel variety produced through 25 generations of selfing. These were treated with colcichine to obtain genotypically matched tetraploid lines. Only the mean and sample size of each cross are reported, and no reciprocal crosses were made. As much as possible, Lundqvist (1966) did attempt to use the same cross direction for the diploids and tetraploids of a given cross, but this was not always possible. The inbred parental lines and F1 were replicated during two experiments in 1956 and 1959 (another attempt was made in 1957 but this failed), whereas the F2 generation was recorded only in 1959. As the 1956 and 1959 replicates generally show a good correlation (see Fig S2A), we take sample-size weighted averages across these two years.

Lundqvist (1966) recorded nine traits for each plant, namely number of spikelets of best-developed ear, seed-setting percentage of best-developed ear, mean kernel weight of best-developed ear, diameter of thickest region of biggest straw, plant height, straw weight, mean and total kernel yield, and number of spikes. Before analyzing the data, we chose “total kernel yield” as the best available proxy for the plant’s overall productivity or fitness.

#### S3.2 Data transformation and interpolation

We transform all the data to find a scaling that generates the strongest correlation between the diploid and tetraploid values, but using only the fixed genotypes (i.e. the inbred parental lines and F1s). In particular, we test a range of values of the scaling parameter *λ* in a Box-Cox transformation, using 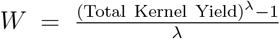. Then, we regress these Box-Cox-transformed diploid and tetraploid values against each other using Standardised Major Axis (SMA) regression using *r*^2^ = Cor^2^(*x* + *y, x* − *y*) to measure goodness of fit. As we only have the line means and not individual-level measurements for these rye data, this transformation ignores the discrepancy between 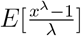 and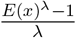. Results reported in Figure S2B-C show that the best-fit *λ* yields a strong correlation between diploid and tetraploid values (*r*^2^ = 0.83), while other *λ* values give a weaker correlation. After the transformation, we use the best-fit SMA regression line to interpolate the fitness value of the missing data point: a tetraploid F1 (black point in Figure S2D).

**Figure S2:**
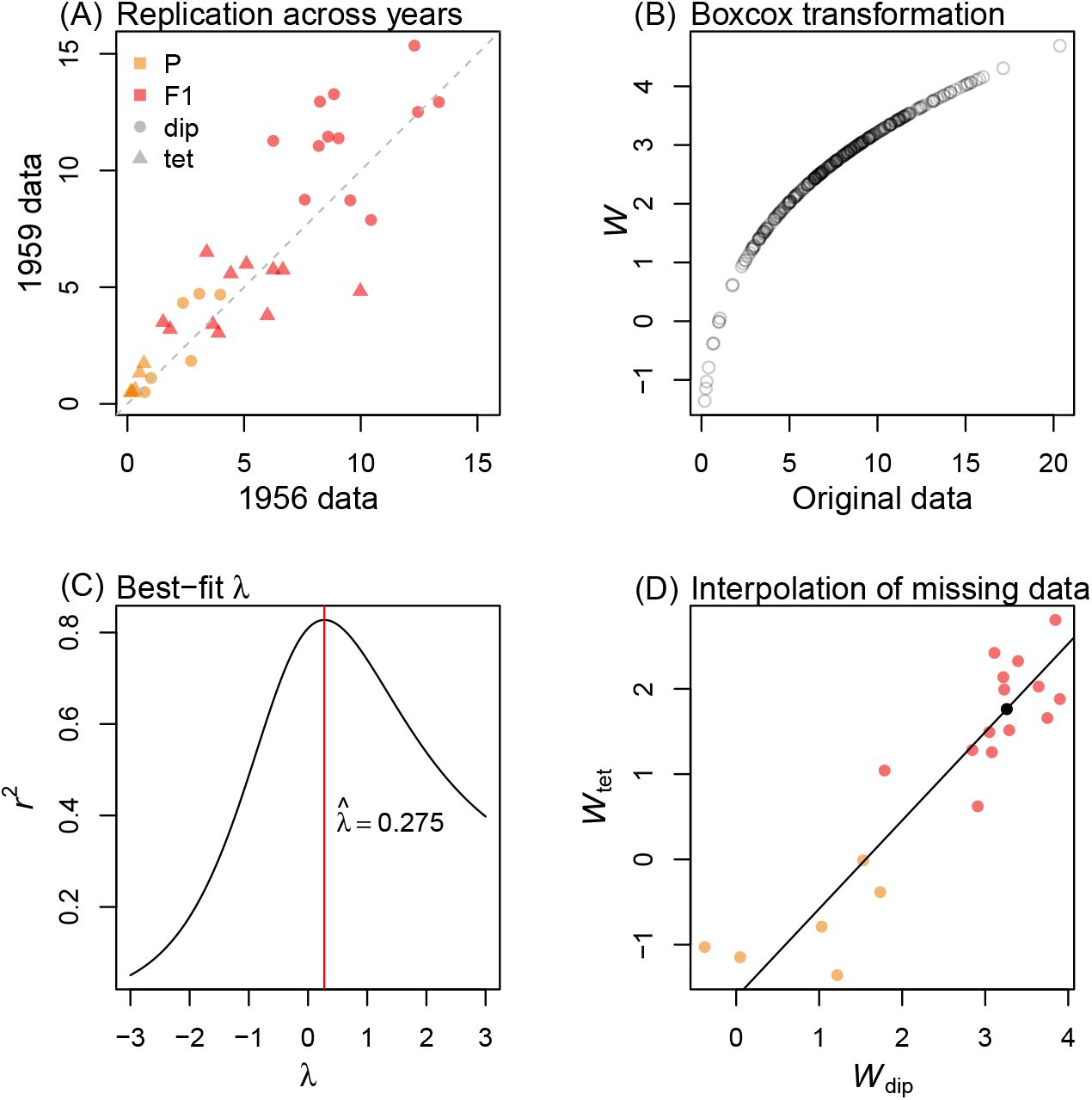
Rye data transformation and interpolation. **(A)** The fitness of diploid and tetraploid (circles and triangles) fixed genotypes (parental lines shown in orange, and F1 in red) is highly correlated between years 1956 and 1959. **(B)** The relationship between the raw and Box-Cox-transformed data. **(C)** The best-fit *λ* = 0.275 (red line) yielded a high correlation in diploid and tetraploid values of fixed genotypes, while other values gave a substantially worse fit. **(D)** With the correlation between the transformed diploid and tetraploid values, we interpolated the missing tetraploid cross (black point) from the best-fit linear model (diagonal line).

**Table S1:** Regression coefficients F2 generation.

| Cross | $P$ | $K$ | $4\langle h_A h_B \rangle$ | $4\langle h_B h_C \rangle$ | $4\sum_{i,j>i}^P \langle h_i h_j \rangle$ | $\langle h_A \rangle \langle h_B \rangle$ | $\langle h_B \rangle \langle h_C \rangle$ | $\sum_{i,j>i}^P \langle h_i \rangle \langle h_j \rangle$ |
| --- | --- | --- | --- | --- | --- | --- | --- | --- |
| AB × AB | 2 | 2 | $\frac{1}{2}$ | — | $\frac{6}{12}$ | $\frac{1}{4}$ | — | $\frac{4}{16}$ |
| | | 4 | $\frac{5}{6}$ | — | $\frac{10}{12}$ | $\frac{1}{4}$ | — | $\frac{4}{16}$ |
| AB × AC | 3 | 2 | $\frac{1}{4}$ | $\frac{1}{4}$ | $\frac{9}{12}$ | $\frac{1}{8}$ | $\frac{1}{16}$ | $\frac{5}{16}$ |
| | | 4 | $\frac{5}{12}$ | $\frac{1}{4}$ | $\frac{13}{12}$ | $\frac{1}{8}$ | $\frac{1}{16}$ | $\frac{5}{16}$ |
| AB × CD | 4 | 2 | 0 | $\frac{1}{4}$ | $\frac{12}{12}$ | $\frac{1}{16}$ | $\frac{1}{16}$ | $\frac{6}{16}$ |
| | | 4 | $\frac{1}{6}$ | $\frac{1}{4}$ | $\frac{16}{12}$ | $\frac{1}{16}$ | $\frac{1}{16}$ | $\frac{6}{16}$ |

#### S3.3 Model fitting

To assess the suitability of our model to explain the patterns in this dataset, we fit our model to the full dataset. As our regression coefficients, we use the expected ancestry proportions and heterozygosities following from eq. 16. These are listed in Table S1 for the F2 generation. Equations 17 and 18 follow from averaging across all combinations of parental pairing.

From Figure 2, we found that the best-fit linear regression of the transformed diploid and tetraploid values has a slope close to 1 and a negative intercept. This suggests that we can account for the effects of tetraploidization by using the same set of *M*_*ij*_ and *I*_*ij*_ values for the two ploidies, but with a constant tetraploidy cost (denoted *tet*). To verify this, we compare models using the corrected Akaike Information Criterion (AICc), which is suitable for parameter-rich models (Burnham and Anderson, 2002). Indeed our model has a lower (i.e. better) AICc value than a model with ploidy-specific *M*_*ij*_ and *I*_*ij*_ values (AICc of 385.33 vs 394.33; see top two rows of Table S2). As Figure 2C revealed, one qualitative failure of the model is the shift of the F2 along the diagonal line, possibly caused by a maternal effect. Adding a maternal effect acting on F2 further decreases the AICc (last row of Table S2). Figure S3 confirms the adequacy of this model for the rye data data, giving a strong correlation of fitted to observed values (panel A), high goodness-of-fit relative to permutations of the ancestry labels (panel B), and constant and approximately normally distributed residuals (panels C and D). Moreover, this model produces sensible parameter estimates. The estimates of *M*_*ij*_ are all large and significantly positive with slight variation among pairs of lines, while *I*_*ij*_ estimates are slightly negative but mostly not significantly different from zero (Figure S3E). This is exactly predicted when parental lines diverged under random drift with possibly very weak stabilizing selection. As anticipated, we also find a significant fitness cost associated with tetraploidy, and fitness boost of the maternal effect acting on the F2 generation (Figure S3F). Figure S4 shows that the model fit for diploid and tetraploid F2 breakdown is substantially improved after accounting for this maternal effect (compare red and black line).

**Figure S3:**
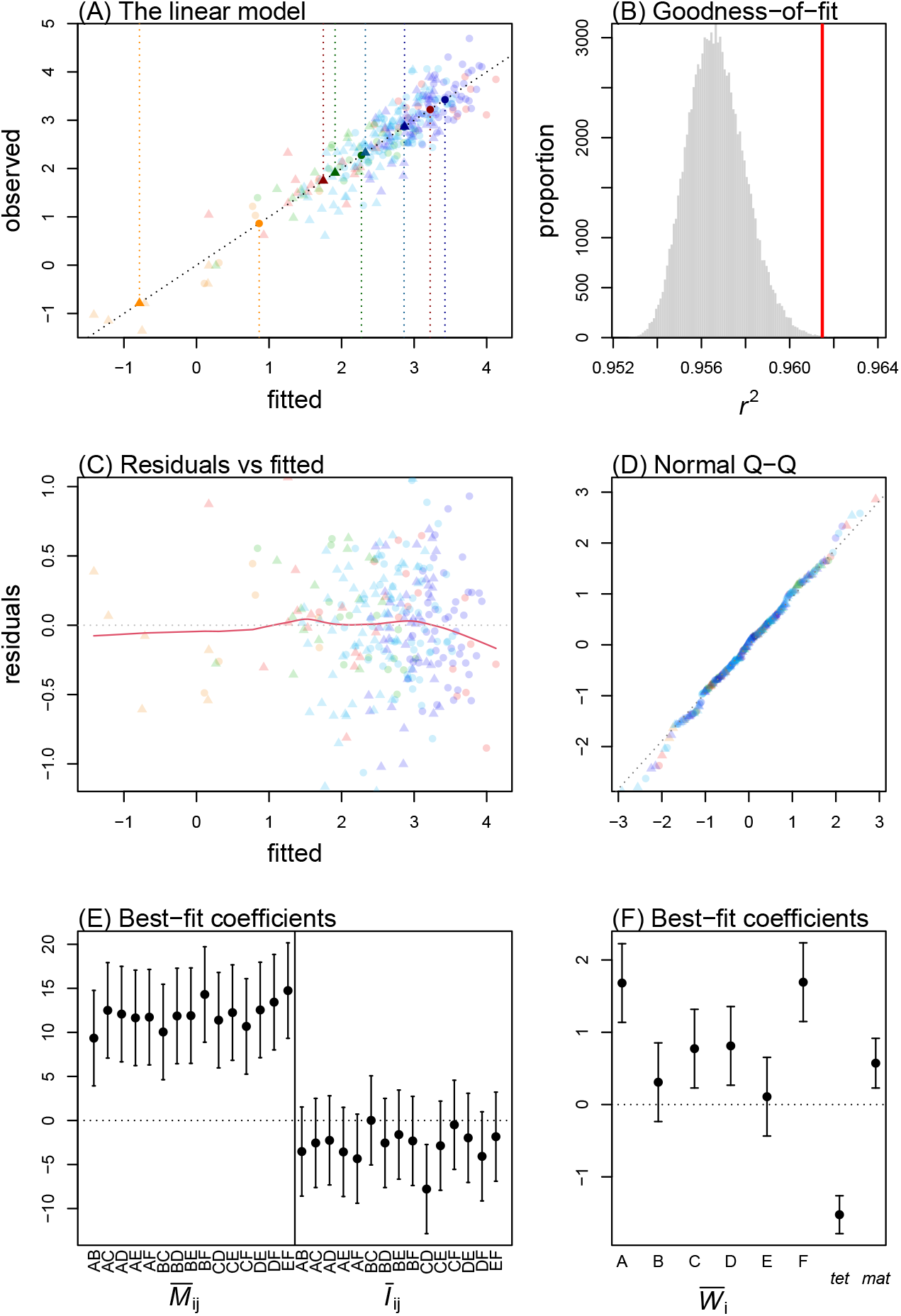
Model fit to rye data. **(A)** The preferred model gives a good fit to the rye data. Faint circles and triangles show the fitted and observed values of each diploid and tetraploid cross. The mean for each cross type is indicated with a clear point and vertical dotted line. **(B)** The *r*^2^ correlation coefficient of the preferred model (red line) lies outside the distribution obtained from fitting the same model after shuffling the ancestry labels (grey histogram). **(C)** The residuals of the model (y-axis) are evenly dispersed across the fitted values (x-axis). **(D)** Normal Quantile-quantile plot shows that model residuals are approximately normally distributed (falling on the diagonal). **(E)-(F)** The parameter estimates under the preferred model indicate that *M*_*ij*_ is similar for most parental pairs, epistasis is on average slightly negative, parental fitnesses vary significantly, and being tetraploid and having a1n3inbred mother both significantly reduce fitness.

**Figure S4:**
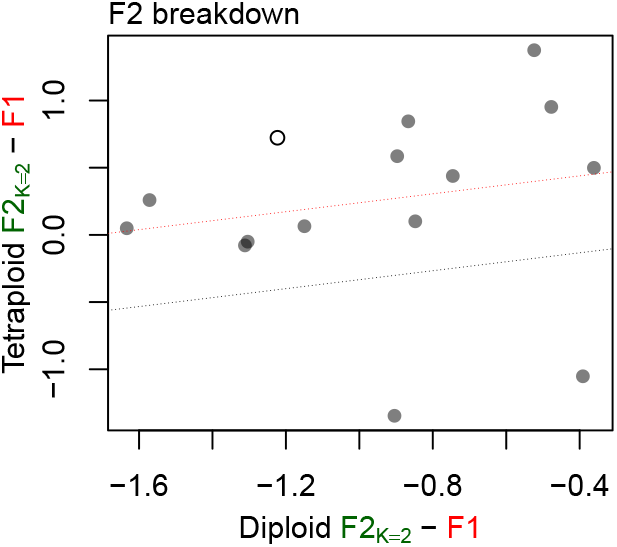
F2 breakdown in rye. All two-parent F2 have lower fitness than F1 in diploids, but not in tetraploids. The red and grey dotted lines show the expected relationship between the degree of F2 breakdown in diploids and tetraploids, following from eq. 14, with and without the estimated maternal effect (slope=1/3, intercept=0.57 or 0). The open point shows the interpolated tetraploid value (see Fig. S2D).

**Table S2:**
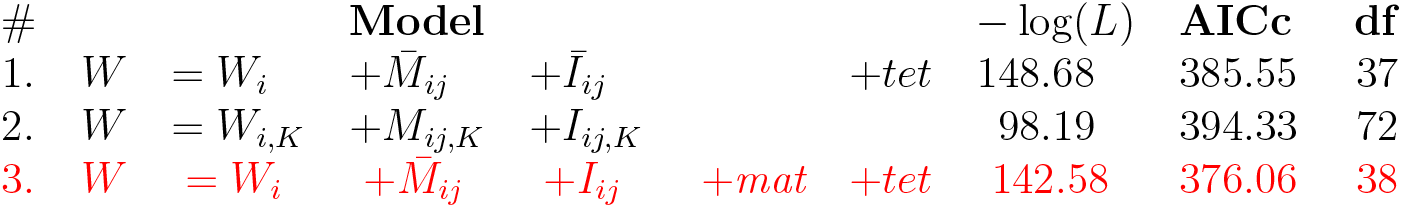
Model comparison for rye data.

**Figure S5:**
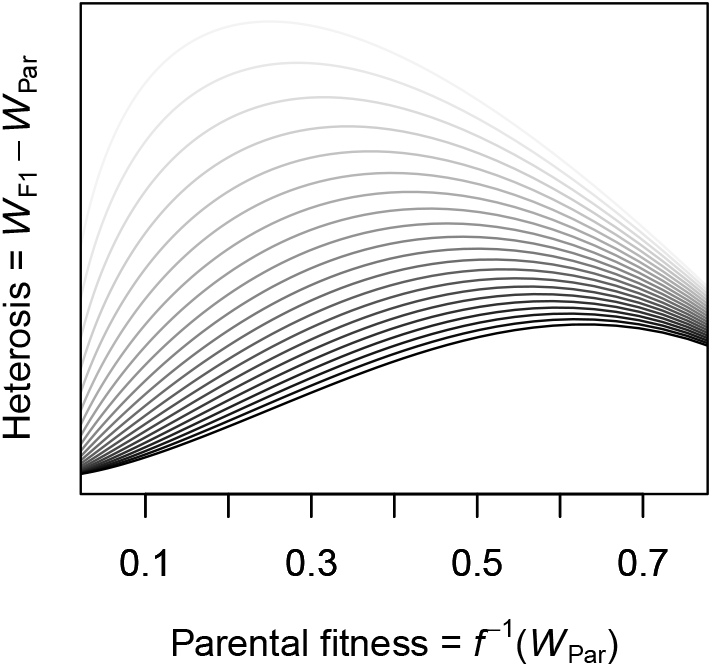
Change in heterosis with parental improvement depends on transformation. Heterosis may increase, decrease or remain constant as parental fitness increases, depending on the transformation used. Grey scale indicates *λ* value used in Box-Cox-transformation, from 1 (light grey) to 3 (black).

## S4 Appendix 4

### Heterosis and parental improvement

Birchler (2013) suggested that the dominance theory of heterosis implies that the level of heterosis should decrease as parental lines increase in fitness (e.g. through purging their deleterious mutations). This pattern was indeed observed by Duvick et al. (2005) in maize for F1 hybrids between lines stemming from different decades of vast yield improvement (although see Troyer and Wellin (2009)). However, the relationship between the level of heterosis and parental fitness depends on the transformation of the data. Figure S5 illustrates increasing parental fitness along the x-axis, and the predicted level of heterosis under our model on the y-axis, for different Box-Cox-transformations i.e. values of *λ* (compare light to dark curves). In all cases, the level of heterosis first increases faster than parental fitness, reaches a maximum, and then decreases with parental fitness. The latter phase is described verbally by Birchler (2013). Yet, the range of parental fitness where each of these regimes occurs depends on the data transformation. Hence, data of this kind do not provide a simple proof for or against the dominance theory as an explanation for heterosis.

